# Sanjeevani: A manually curated anti-cancerous phytochemical database integrated with tools

**DOI:** 10.64898/2026.06.15.732344

**Authors:** Vicky Jha, Ronak Jha, Sakshi Shukla, Shardul Shingan, Gourab Das

**Author notes:** Corresponding author: Gourab Das.

## Abstract

**Background:** Cancer continues to pose a massive global health burden. While plant-derived phytochemicals offer promising therapeutic leads, existing natural product databases often lack cancer specificity, dataset downloadability, and integrated screening tools.

**Methods:** We developed Sanjeevani, an integrative web platform cataloguing 4,823 curated anticancer phytochemicals. Using a balanced dataset of 9,646 molecules, we trained Support Vector Machine (SVM), Random Forest, and K-Nearest Neighbours classifiers using a hybrid feature representation of RDKit descriptors and 2048-bit ECFP4 fingerprints. The platform also integrates AutoDock Vina for web-based molecular docking for binding affinity, poses prediction and ADMET-AI for pharmacokinetics estimation.

**Results:** The SVM model demonstrated the strongest predictive capability, achieving a top test accuracy of 0.966 and a ROC-AUC of 0.992. Benchmarking across five docking tools confirmed that AutoDock Vina successfully balanced computational automation with literature-consistent binding affinity replication. The final architecture provides rapid interactive 2D/3D visualizations integrated with downstream analysis tools.

**Conclusion:** Sanjeevani provides an open-access, one-stop pipeline that bridges the gap between raw natural product data and actionable computational screening, accelerating natural product-based oncology drug discovery.

**GRAPHICAL ABSTRACT:** 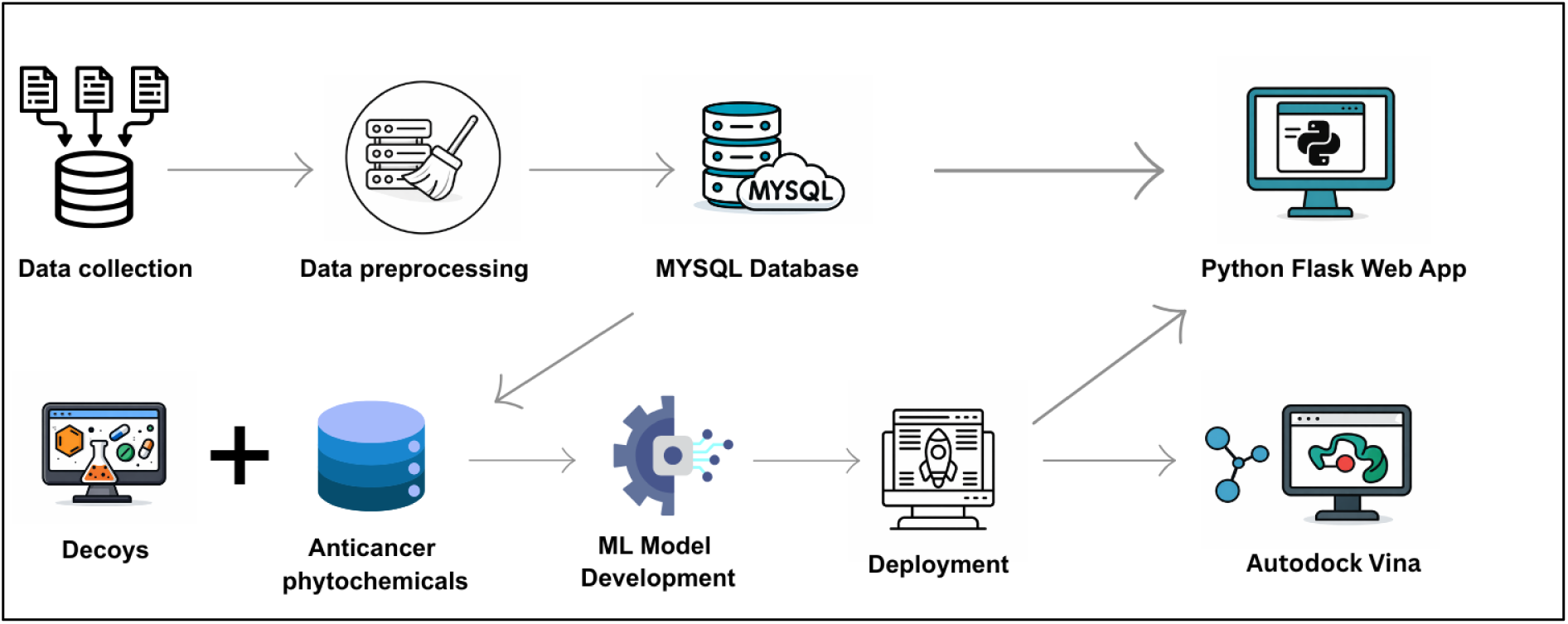

## INTRODUCTION

Cancer remains a major global health challenge, causing an estimated 20 million new cases and over 10.3 million deaths in 2025 alone. Despite enormous progress in molecular biology, the global burden of cancer is on the rise, creating significant clinical and economic challenges. Hallmarks of cancer including genomic instability, immune evasion, and metastasis continue to limit therapeutic success and contribute to cancer’s status as a “modern-day poison”[1]. Advances in computational and molecular technologies have accelerated the search for targeted interventions and new anticancer compounds. Phytochemicals bioactive compounds derived from plants have garnered growing attention as promising anticancer agents. These molecules exhibit diverse chemical structures and modulate key cellular pathways implicated in cancer development, such as proliferation, apoptosis, angiogenesis, and metastasis. Plant-derived anticancer agents, including paclitaxel, vincristine, and camptothecin, have played central roles in chemotherapy protocols while offering favorable toxicity profiles compared to synthetic drugs [2]. Because they are often optimized by evolutionary processes, phytochemicals can serve as efficient lead compounds for drug discovery [3]. Drug discovery efforts rely increasingly on large-scale computational screening, including artificial intelligence and machine learning, to predict biological activity and optimize lead identification [4]. Nevertheless, research is hampered by major limitations in existing natural product databases [5]. Several databases have been developed to catalog phytochemicals with potential anticancer activity. Among them, NPACT (Naturally Occurring Plant-based Anti-Cancer Compound Activity-Target database) provides a curated list of compounds experimentally validated against cancer cell lines and targets [6]. CancerHSP focuses on herbal secondary metabolites with anticancer properties [7], while Dr. Duke’s Phytochemical and Ethnobotanical Database offers valuable ethnobotanical insights and chemical profiling of medicinal plants [8]. SANCDB (South African Natural Compounds Database) provides a referenced collection of natural compounds from South African biodiversity with structural formats and analog information to support drug discovery [9]. ANPDB (African Natural Products Database) integrates natural compounds with spectral, pharmacokinetic, and ethnobotanical data to enable cheminformatics and AI-assisted drug exploration [10]. BRNPDB (Brazilian Natural Products Database) combines chemical, biological, and geographic information on Brazilian natural products to promote biodiversity-based pharmacological research [11]. AVPCD (Antiviral Phytochemicals Database) focuses on plant-derived antiviral compounds and provides a platform linking phytochemical, disease, and structural information for therapeutic development [12]. However, these databases exhibit key limitations. First, most are not cancer-specific. Second, database downloadability is severely restricted; users may only access lists of compound names rather than complete datasets linking compounds to bioactivity [5]. Third, very few databases provide downloadable 2D and 3D chemical structures, which greatly limits their utility for molecular modeling and docking studies which is a essential steps in computer-aided drug design [5]. Integrated prediction and docking tools, allowing researchers to virtually screen and evaluate phytochemical interaction with human protein targets, are typically absent [5, 13]. To address these gaps, the Sanjevani database provides the largest curated collection of anticancer phytochemicals. This resource curates 4823 unique anticancer phytochemical compounds along with essential identifiers (e.g., Cid, InChIKey, Molecular weight, Smiles and Biological activity), downloadable 2D/3D structures, and embedded docking tool, bioactivity and ADMET prediction. For the first time, researchers can access a single, user-friendly platform that enables compound exploration, virtual screening, and molecular docking, as well as reliable machine-learning based anticancer predictions. The primary goal of Sanjevani is to accelerate natural product based cancer research by providing the global research community biomedical scientists, pharmacologists, computational biologists, and drug developers with an easy-to-use, integrative, one-stop platform for anticancer phytochemicals. This resource empowers users to conduct efficient in silico investigations, bridging discovery and translational application while overcoming bottlenecks in data accessibility, structure availability, and predictive analytics. The overall workflow used for the construction of the Sanjevani database and integration of machine learning, docking, and ADMET prediction modules is illustrated in **Supplementary Figure S1A**.

## Materials and methods

### Phytochemical Data Retrieval

Information was collected from various public repositories of both ethnobotanical and natural product databases in order to assemble a comprehensive dataset of phytochemicals with experimentally validated anticancer activity. Sources reporting biological or organ-specific anticancer assays were prioritized in the retrieval strategy to ensure relevance and diversity across compound classes. Data extraction was performed through a combination of bulk downloads and automated web scraping pipelines implemented in Python. Repositories such as Dr. Duke’s Phytochemical and Ethnobotanical Databases, NPACT, and CancerHSP served as the principal resources due to their extensive coverage of plant-derived compounds with annotated anticancer bioactivities. Additional regional and thematic repositories including SANCDB, BrNPDB, ANPDB, and AVPCD were incorporated to capture geographically diverse natural products and cell line specific bioactivity data. After combining data from all sources, these compounds form a diverse and well-curated library of plant derived molecules with reported anticancer properties providing a strong foundation for future computational analyses, predictive modeling, and bioinformatics-based drug discovery.

**Table 1:**
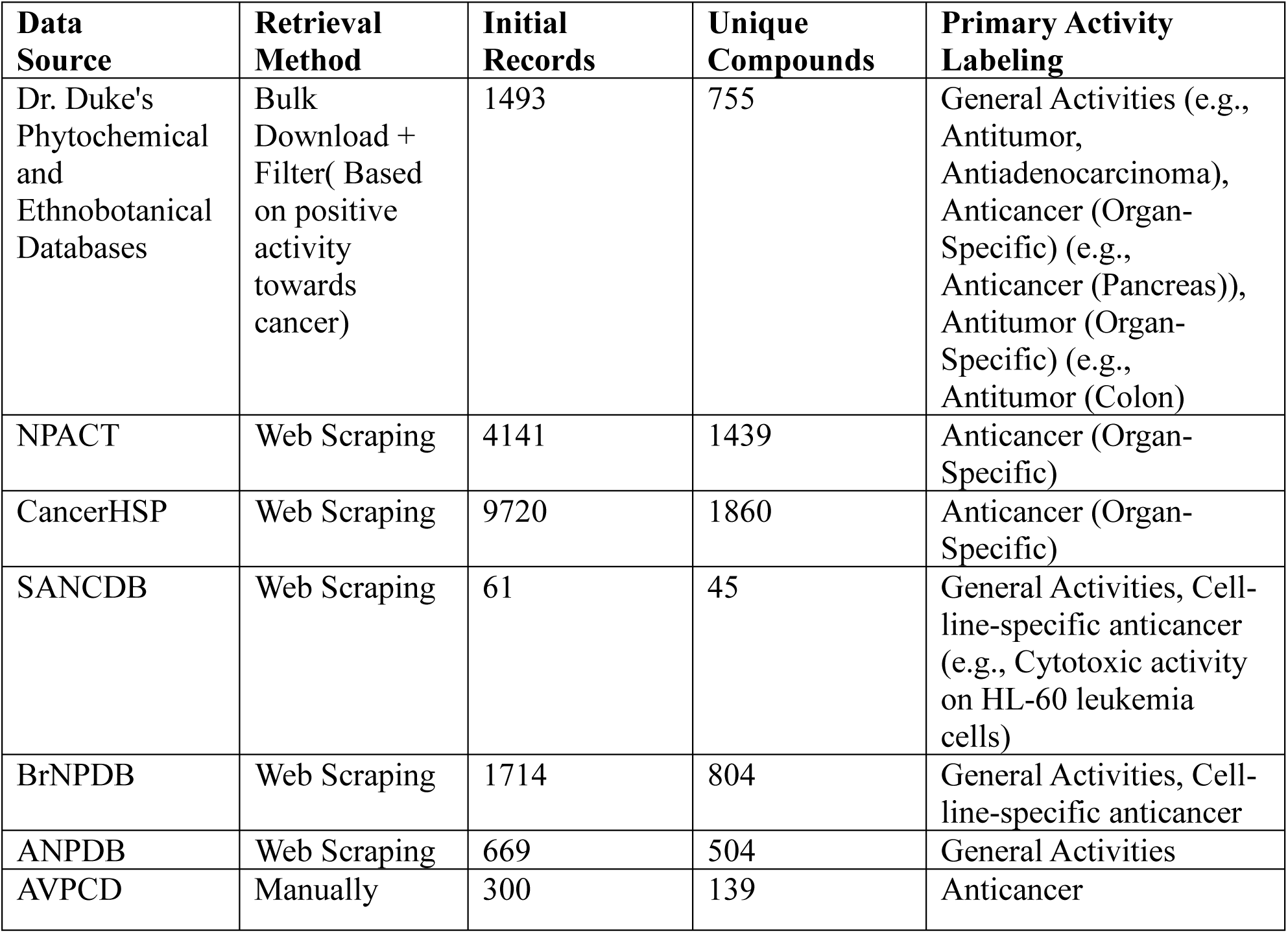
Summary of phytochemical data sources and compound extraction workflow for the Sanjevani database.

### Data Enrichment and Standardization

Following the initial compilation of phytochemicals, compound standardization and data enrichment were performed to ensure uniformity across all integrated sources. The InChIKey was used as the primary query parameter for retrieving additional molecular information from the PubChem REST API, ensuring exact structural matches across databases. This approach avoided inconsistencies that often arise when compound names differ between sources, thereby improving the accuracy of data retrieval. Each compound was enriched with key molecular descriptors, including the PubChem Compound ID (CID), molecular weight, and canonical SMILES string. This enrichment ensured that all entries were represented with standardized, machine-readable chemical identifiers suitable for cheminformatics and bioinformatics analyses. Compounds lacking corresponding entries in PubChem were excluded to maintain dataset integrity and reliability. After enrichment, datasets from all repositories were merged into a single, harmonized collection, which consolidated multiple records corresponding to the same compound from different sources. This refinement reduced the dataset from an initial pool of 18,100 entries to 11,194 compounds containing complete structural and molecular information. While initial deduplication removed structurally identical compounds across databases, entries with the same molecular structure but distinct biological activities were intentionally retained to preserve activity diversity. To handle this overlap without losing biologically relevant information, the dataset was further standardized and reorganized using the InChIKey as the unique compound identifier. The data were then normalized following Third Normal Form (3NF) principles through Python-based preprocessing. This process separated compound structures, activity descriptions, and their relationships into independent yet interconnected tables, ensuring that each biological activity could be accurately linked to its corresponding compound without duplication. By applying this structured normalization, the final harmonized dataset contained 4,823 unique phytochemicals, each associated with one or more validated biological activities. This approach ensured both the chemical integrity and biological richness of the dataset while optimizing it for efficient querying and downstream computational analysis.

### Structural Data Retrieval

Two-dimensional (2D) and three-dimensional (3D) structural data for the phytochemical compounds were retrieved in SDF format using the PubChem REST API.

### Database design

The backend architecture of Sanjeevani was implemented using Flask, a Python-based web framework that supports lightweight and modular integration of database-driven analytical applications. The application was deployed with a web-accessible interface that enables interaction between the user-facing platform and the underlying computational modules through server-side request handling and route-based execution. The data layer of the platform was organized using a MySQL relational database, designed to enable efficient storage, retrieval, and querying of curated phytochemical and anticancer activity records. The database schema was structured to support rapid access to compound specific information and associated biological annotations while maintaining referential consistency across interconnected tables.

**Supplementary Figure S1A:**
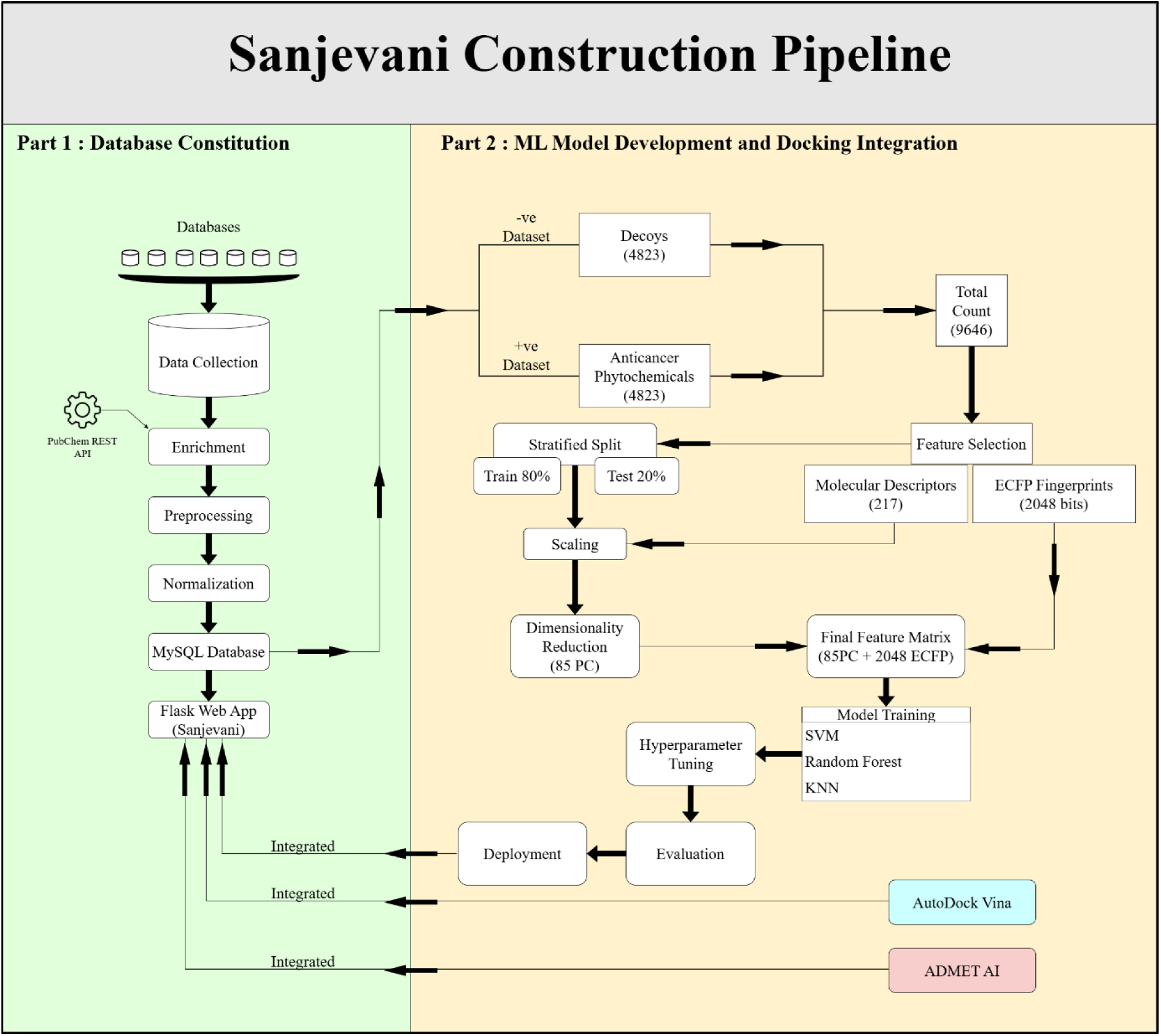
**Part 1:** Phytochemical data were collected from multiple chemical databases and enriched using the PubChem REST API. The curated data underwent preprocessing and normalization before being stored in a MySQL database and deployed through a Flask-based web application named Sanjeevani. **Part 2:** A balanced dataset of anticancer phytochemicals and decoy compounds was then prepared for machine learning analysis. Molecular descriptors and ECFP fingerprints were generated, followed by stratified splitting, scaling, and PCA-based dimensionality reduction. Machine learning models were trained and optimized, and the final system was integrated with docking using AutoDock Vina and pharmacokinetic prediction using ADMET AI within the Sanjeevani platform. **(Note this will go to supplementary table)**

### Access and Web Interface

The Sanjevani platform is publicly accessible through its dedicated web portal at https://sanjeevani.actrec.gov.in/. The web interface provides an intuitive and user-friendly environment for browsing, visualizing, and analyzing phytochemical compounds with reported anticancer activity. Upon accessing the Sanjevani homepage, users are introduced to an overview of the database, including its data sources, features, and integrated computational tools. The portal is organized into modular sections accessible via the top navigation bar Home, Browse, Tools, Contact, and Download each designed to streamline specific research tasks.

**Figure 1.**
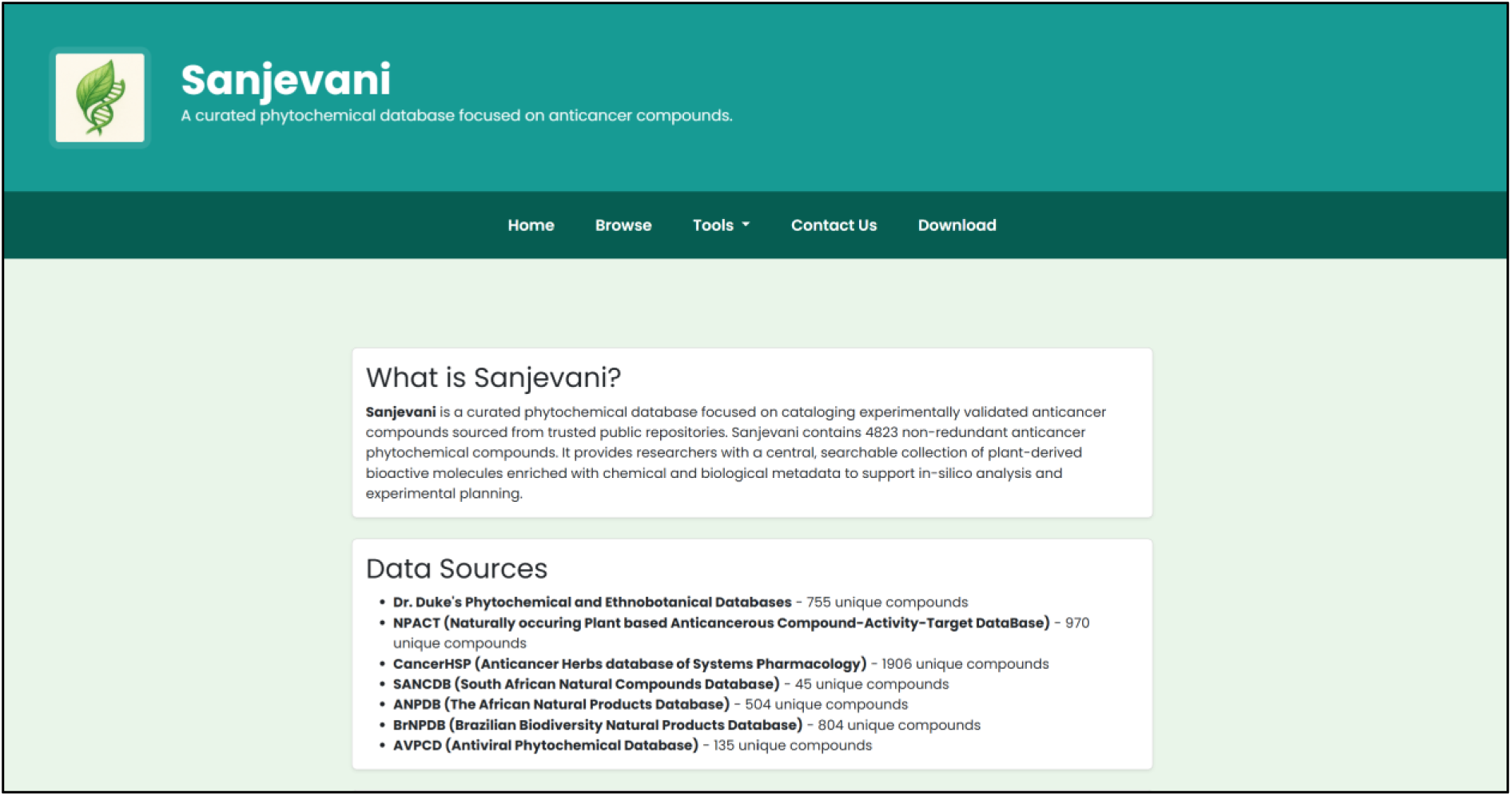
Sanjevani Homepage, Navigation Bar and its information

### Browse Page

The Browse section presents the curated phytochemical dataset in a searchable and sortable datatable implemented with Flask and JavaScript-based live filtering. Users can perform keyword searches by compound name or biological activity, and dynamically control the number of records displayed (10, 50, or 100 entries per page). Each table entry lists the compound name, PubChem CID, InChIKey, molecular weight, SMILES string, and biological activity, accompanied by a “More” button that links to the Compound Detail Page. The Compound Detail Page provides complete molecular and activity information for the selected compound. It integrates 2D and 3D molecular visualizations using SmilesDrawer and 3Dmol.js, respectively. The 3D viewer offers multiple interactive visualization modes ball-and-stick, stick, wireframe, and spacefill along with optional hydrogen toggling and automatic spin animation. A built-in download panel allows users to export molecular data as 2D PNG images or 2D/3D SDF files for Molecular docking.

**Fig 2:**
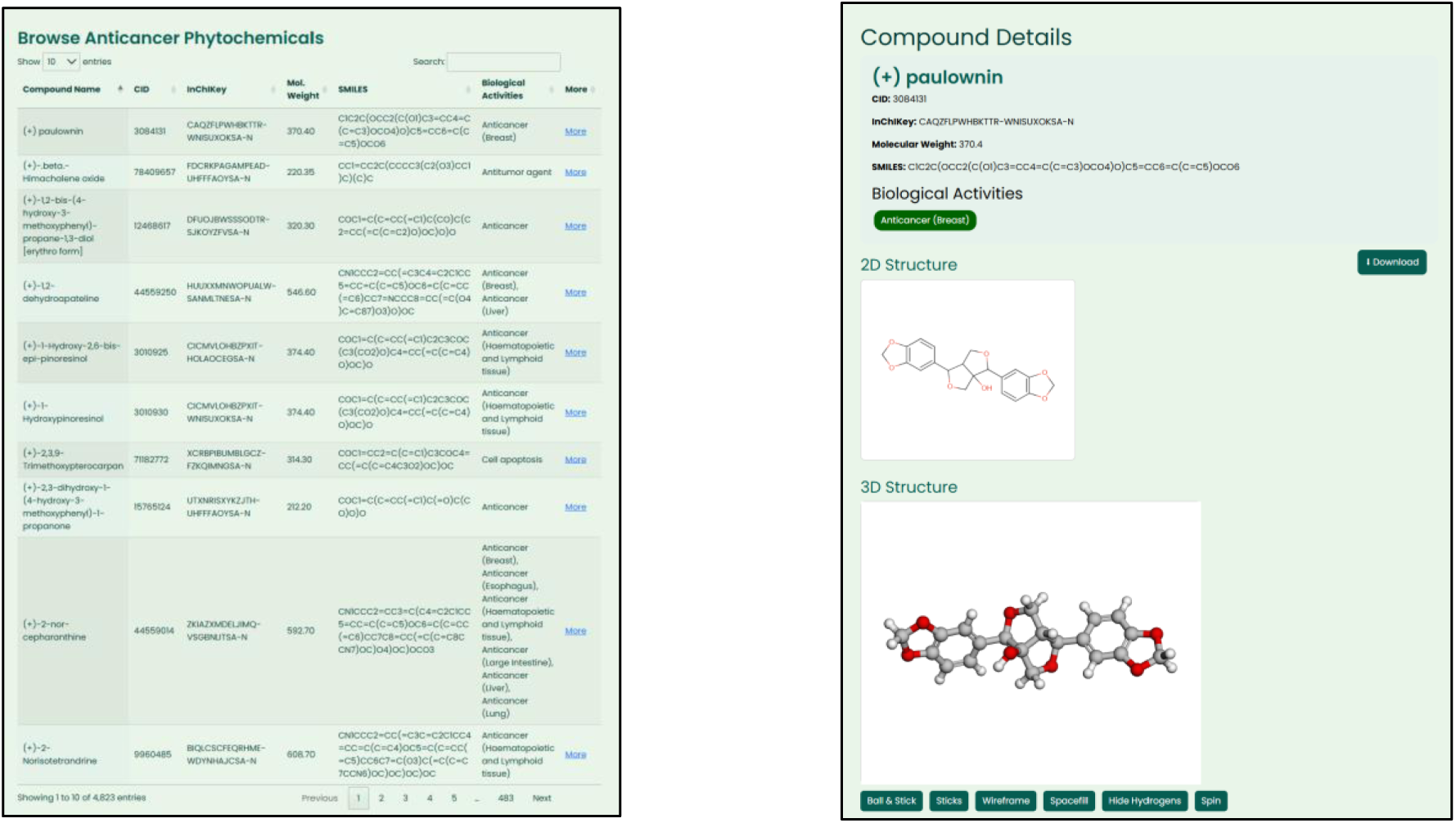
Browse Page

### Tools Page

The Tools section hosts two interactive modules designed to support cheminformatics and computational screening tasks:

### Docking Tool

This module integrates AutoDock Vina (v1.1.2) for automated receptor-ligand docking. Users can upload receptor and ligand structures in PDBQT format, specify docking box coordinates (center and dimensions), and adjust parameters such as exhaustiveness, number of binding modes, and CPU usage. The results, including binding affinities and ranked docking poses, are displayed on the same page. Users can visualize docking complexes in real time via 3Dmol.js, adjust molecular representations (cartoon, stick, surface, wireframe), and download the output and log files for further analysis.

**Fig 3:**
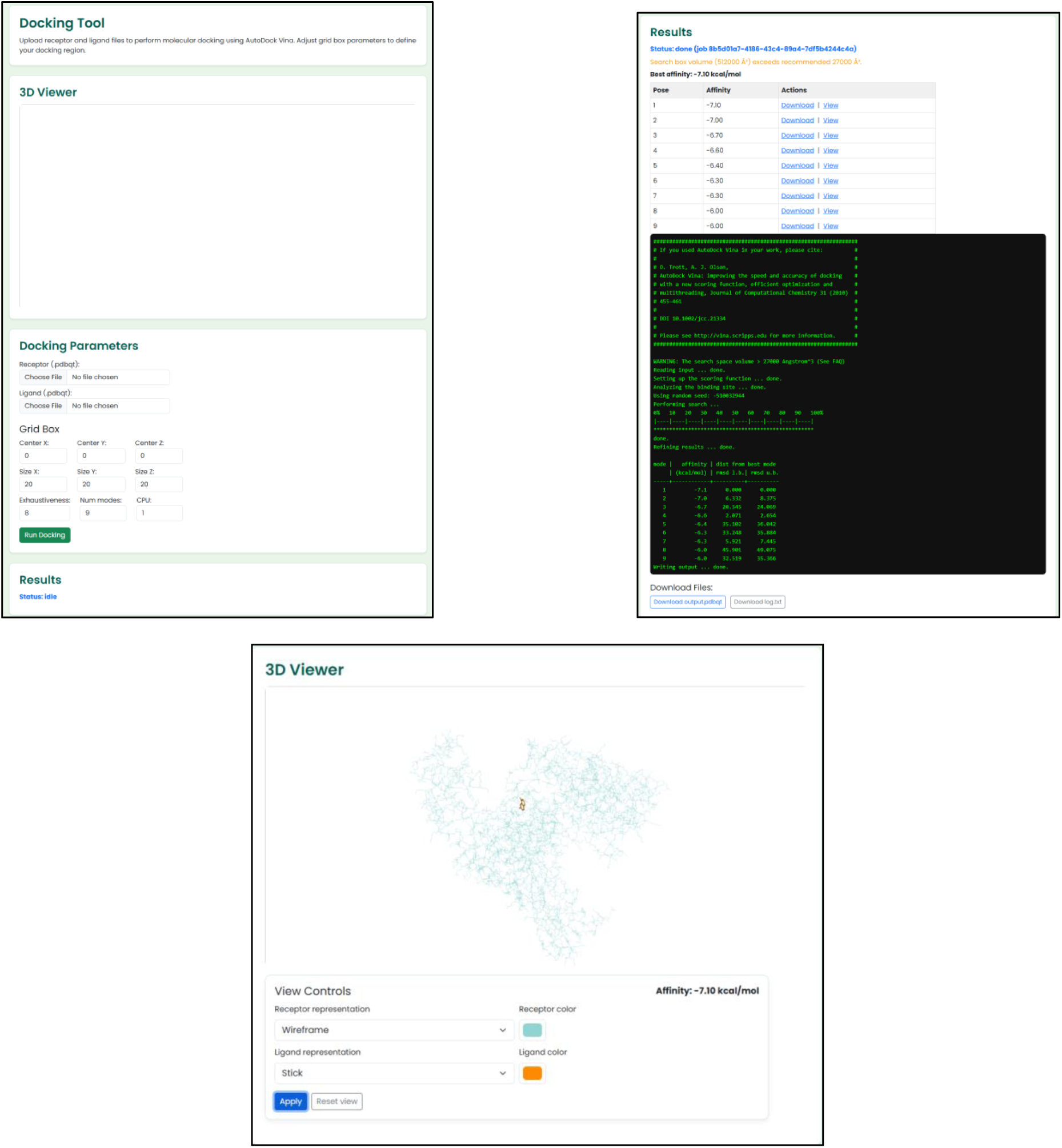
Docking Tool Page

### Prediction Tool

The anticancer activity prediction module implements a Support Vector Machine, Random Forest classifier and K-Nearest Neighbour trained on 4,823 curated phytochemicals. Users input a SMILES string and select the model, and the model outputs a binary classification “Anticancer” or “Non-anticancer” along with prediction probability. The backend employs RDKit for feature generation and the scikit-learn pipeline (serialized via Joblib) for inference.

**Fig 4:**
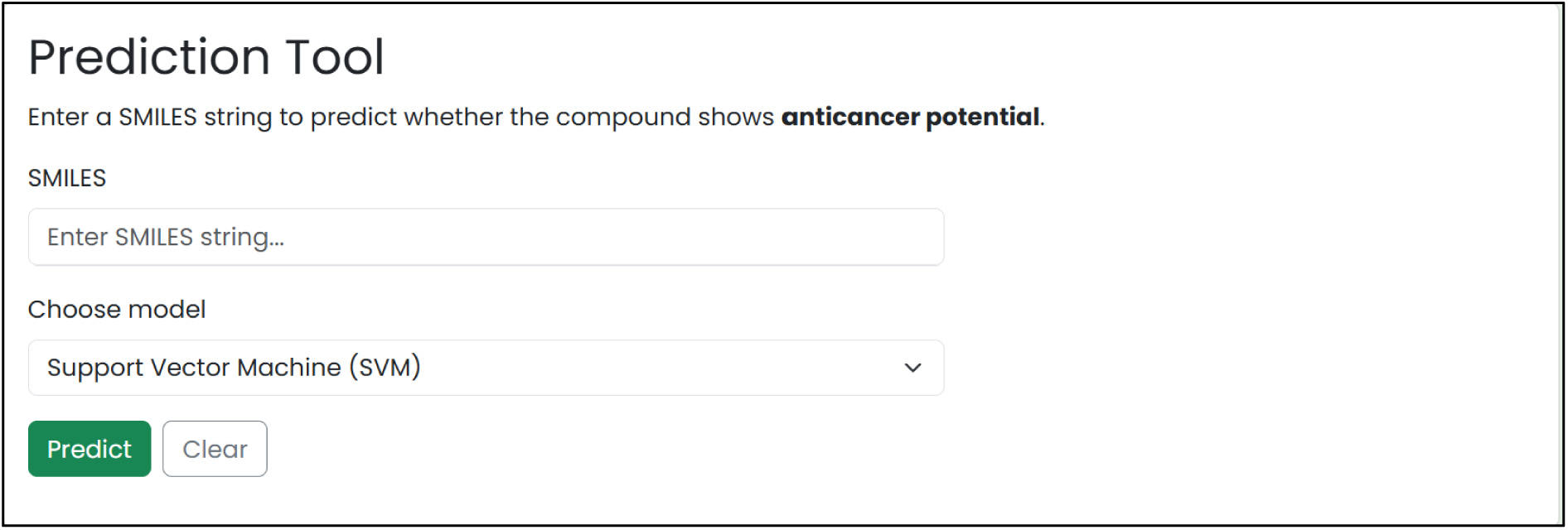
Prediction Tool Page

### ADMET Tool

The ADMET-AI module allows users to input a compound’s SMILES string and instantly obtain predicted ADMET and drug-likeness properties, along with physicochemical descriptors and percentile-based comparisons to approved drugs, facilitating rapid evaluation and prioritization of candidate molecules.

**Fig 5:**
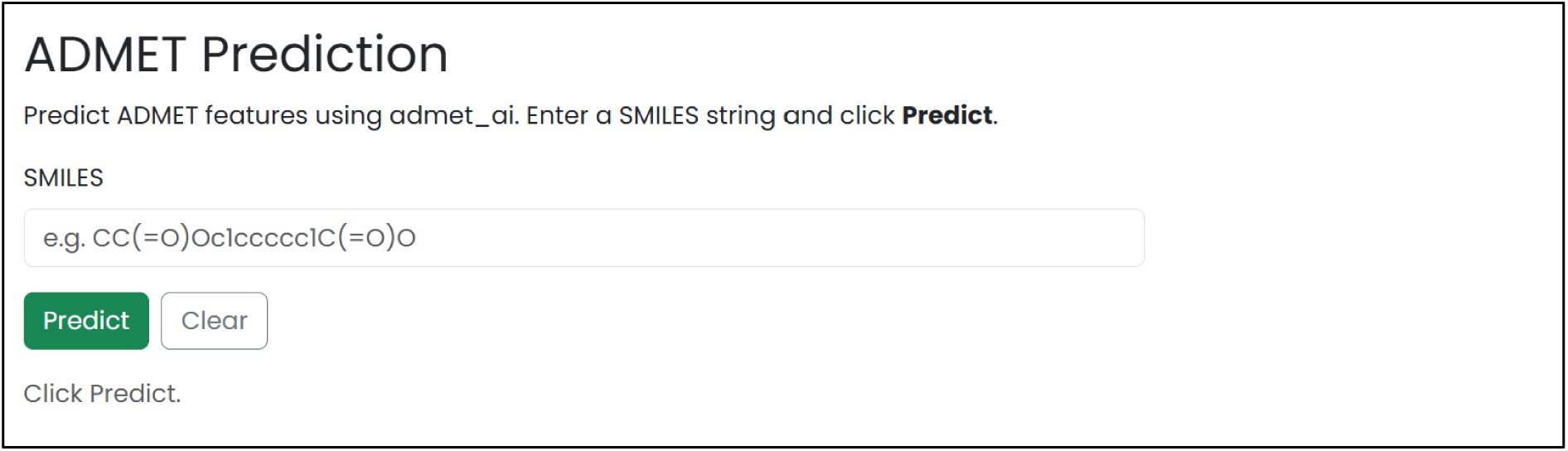
ADMET Prediction Tool Page

### Download and Contact

The Download page allows users to obtain the complete Sanjevani dataset, including all compounds and associated biological activity information, in Excel (.xlsx) format for offline or large-scale computational analysis. The Contact page provides the developer’s contact details (email) for inquiries, collaborations, and feedback.

## Results

### Platform Performance Overview

The Sanjevani web application was successfully implemented and validated for stability, responsiveness, and functional integration. All core modules including the data browsing interface, molecular visualization, anticancer prediction model, docking tool and ADMET tool performed as intended without critical errors during execution. The web interface demonstrated smooth rendering of 2D and 3D structures through SmilesDrawer and 3Dmol.js, with average page load times below 2 seconds, confirming the framework’s efficiency for interactive cheminformatics analysis… (**Deployment point with docker, queue management will come here(we will write about the time the docking takes and how many people at one time it supports)…just a tentative paragraph)**

### Data collection and preprocessing for machine learning model generation

A total of 9,646 compounds were used for machine learning model development, consisting of both active and inactive molecules. The data was randomly divided into a test set and a training set in 80:20 ratio. At each of the 10 iterations, 7,716 compounds (80%) were placed into the training set, and 1,930 compounds (20%) were placed into the test set. Stratified sampling was used in order to maintain the original class distribution in each partition. For molecular representation, two types of features were employed, molecular descriptors which in total was 217 and ECFP4 fingerprints which was 2048 bits. The subset of descriptors was only subjected to Principal Component Analysis (PCA). PCA also minimized the initial space of the descriptors to 85 principal components which captured most of the variance of the data. The ECFP4 fingerprints were not reduced in dimensions, hence fine substructural information, which is essential in bioactivity prediction, is preserved. Each compound was then taken and its final feature vector was formed by concatenating the 85 PCA transformed items of the descriptors with the 2048-bit ECFP4 fingerprint to obtain a total feature dimension of 2,133.. The same preprocessing and feature dimensions were used consistently for the Random Forest, Support Vector Machine, and k-Nearest Neighbors models, ensuring uniform input representation across all classifiers.

### Model Generation

In developing the models, three supervised machine-learning models, namely, the Random Forest (RF), Support Vector machine (SVM), and K-Nearest Neighbors (KNN) were implemented using the scikit-learn library in Python. All the models were trained and tested ten times separately. Each of the runs used random division of the data, preprocessing, tuning with the help of GridSearchCV with 10 cross validation, and eventually evaluated on an independent test set. The performance measures derived cross-validated accuracy, training accuracy, test accuracy, ROC-AUC, precision, recall, F1-score, and Matthews correlation coefficient (MCC)) of all ten runs and are summed-up in **Supplementary Table S1.** According to the comparative analysis of the performance in the test-set concerning all their iterations, the Support Vector Machine (SVM) revealed the best predictive power. The most effective model of SVM had the highest test accuracy (0.966), highest values of precision (0.963), recall (0.979), F1-score (0.963), MCC (0.932) and ROC-AUC (0.992), indicating the model have a strong discriminative factor, and balances between active and inactive compounds. Random Forests demonstrated the second best performance, the test accuracy of about 0.958, high ROC-AUC of (0.988), and strong values of MCC (0.917), dramatically demonstrates its stable performance and its capacity to work well with the large dimensional data. The k-Nearest Neighbors classifier was also shown a maximum of 0.954 test and about 0.908 MCC. Overall, the models were ranked in the following manner SVM > RF > KNN using test-set performance metrics. As a result, the optimal RF, SVM, and KNN models, which were the most effective models during the repeated iteration were chosen to serve as the final predictive models and have then been incorporated into the Sanjevani web platform to do the virtual screening and identification of the anti-cancer phytochemicals.

**Table 9:** Performance Evaluation of the SVM, RF and KNN (Test Dataset)

| Performance | Support Vector Machine | Random Forest | K- Nearest Neighbors |
| --- | --- | --- | --- |
| Accuracy | 0.9663 | 0.9585 | 0.9533 |
| Precision | 0.9629 | 0.9529 | 0.9344 |
| Recall | 0.9699 | 0.9647 | 0.9751 |
| MCC | 0.9326 | 0.9171 | 0.9075 |
| AUC | 0.9899 | 0.9883 | 0.9767 |
| F1-score | 0.9664 | 0.9588 | 0.9543 |

**Fig 6:**
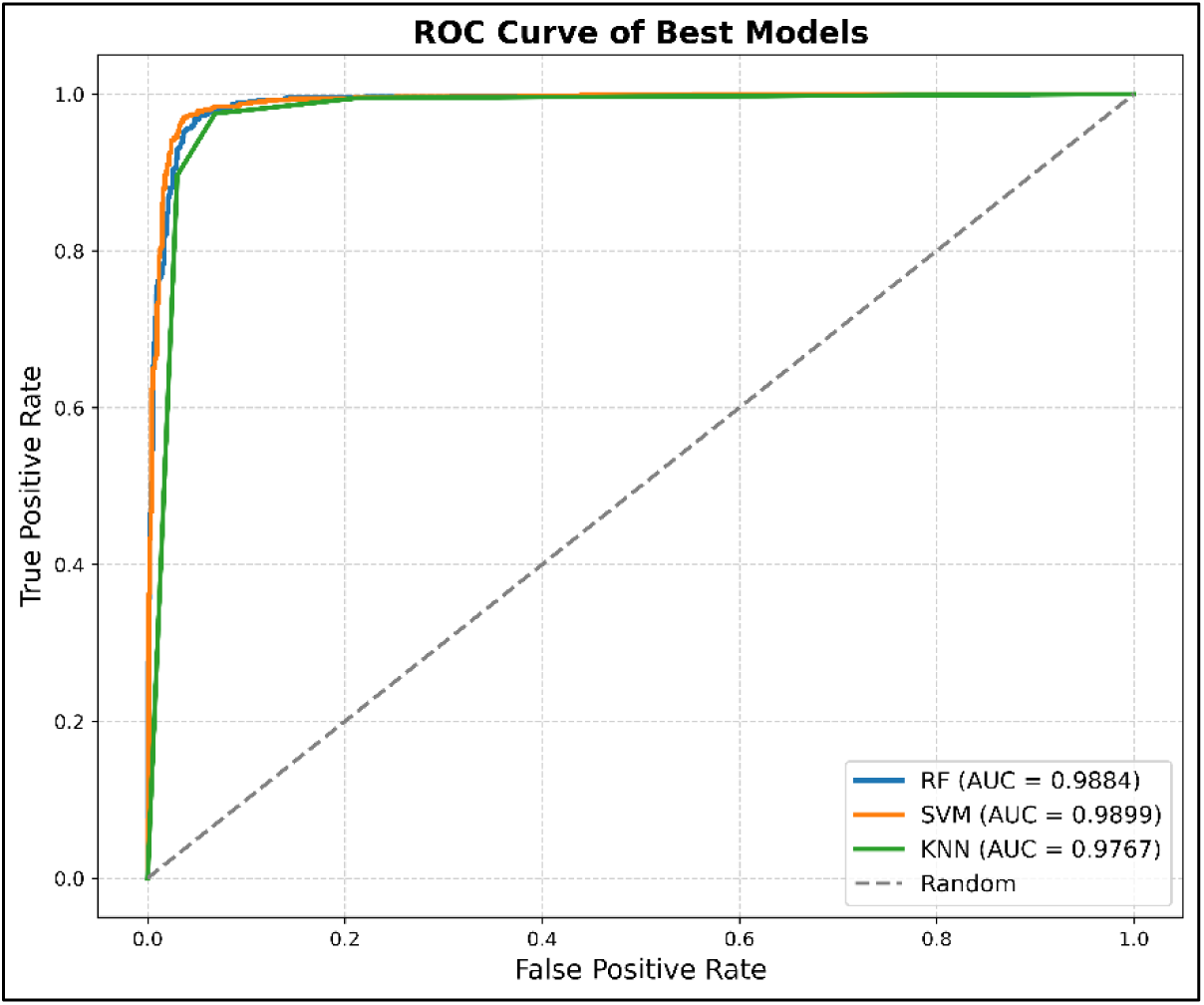
ROC-AUC Curve

### Prediction Models Validation

**Table 10:**
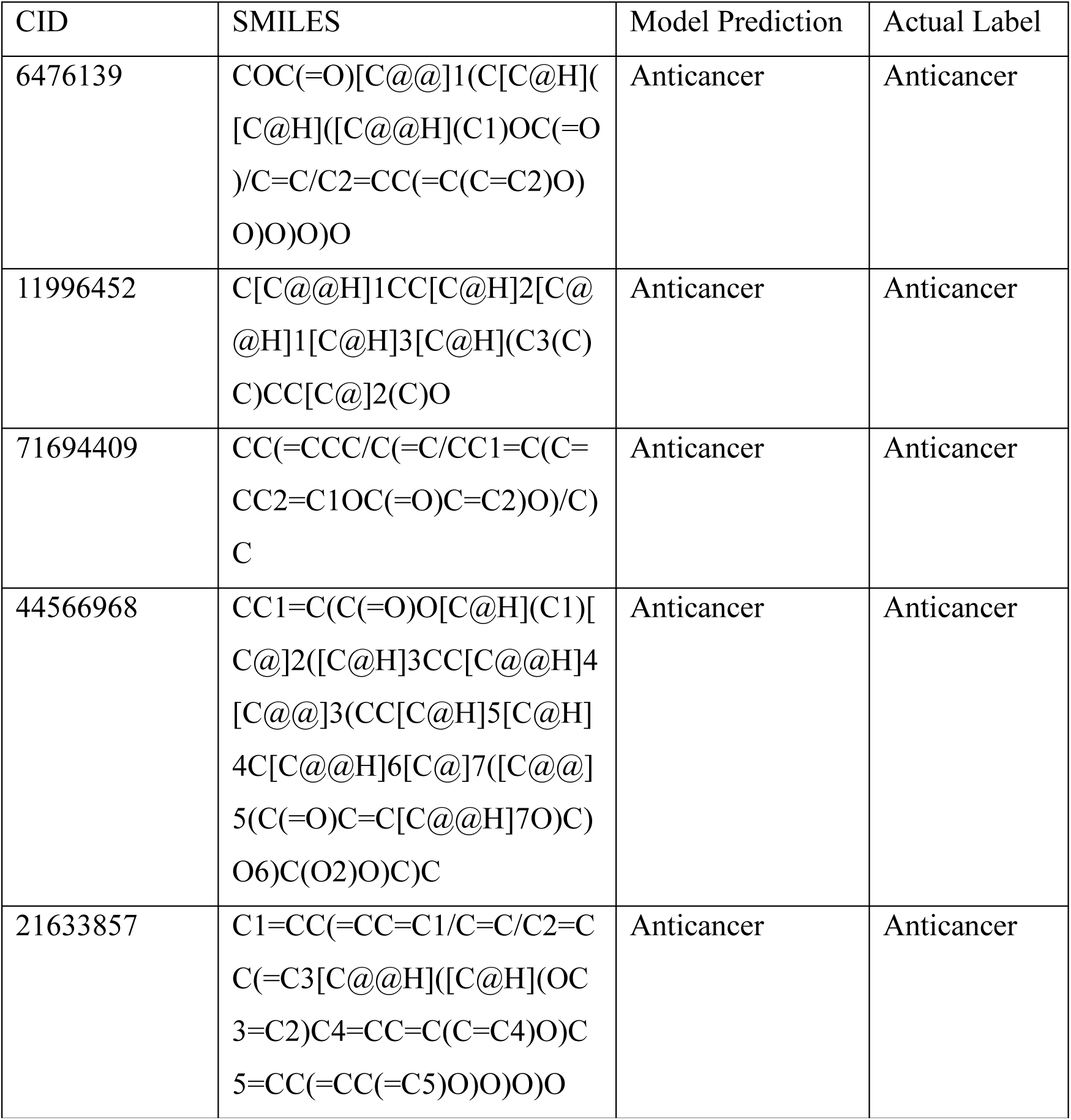
Blind Anticancer prediction of Anticancer phytochemicals [40][41][42][43][44].

**Table 11:** Blind Anticancer prediction of Non-Anticancer phytochemicals [14].

| CID | DECOYS | Model Prediction | Actual Label |
| --- | --- | --- | --- |
| 72521 | O=C(CSc1nc[nH]n1)Nc1ccnc2ccccc12 | Non-Anticancer | Non-Anticancer |
| 24884282 | CCC(COC)Nc1cc(C)nc2c(c3cnc(N(C)C)cc3C)c(C)nn12 | Non-Anticancer | Non-Anticancer |
| 1715097 | COc1cccc(N2CCOCC2)c1 | Non-Anticancer | Non-Anticancer |
| 624491 | Cc1c(CC(=O)NN)c2ccccc2n1c1ccccc1 | Non-Anticancer | Non-Anticancer |
| 3084131 | O=C(Cn1ncc2ccsc2c1=O)N1CCN(c2ccc(Cl)cc2)CC1 | Non-Anticancer | Non-Anticancer |

### Docking Tool Benchmarking

To ensure accurate, reproducible, and computationally feasible integration of a docking engine within the Sanjevani web application, five docking platforms UCSF Chimera, Webina, CB-Dock2, SwissDock, and PyRx were systematically benchmarked. The goal was to identify a docking tool that consistently reproduced literature reported binding affinities across a representative panel of anticancer phytochemicals and protein targets, including VEGFR2, HER2, HER3, HCA-II, h11 Beta-HSD1, hCDC25B, BZLF1 gene, 1FLT and HSP90AA1. As presented in Table 12-14, Chimera demonstrated the most reproducible results, with the highest number of binding affinities, matching and closely approximating literature-reported values. Table 15 further summarizes these outcomes, confirming Chimera as the strongest performer in terms of reproducibility and overall affinity accuracy, followed by Webina and PyRx. However, despite its superior docking precision, Chimera’s heavy graphical user interface (GUI) and desktop dependent architecture is unsuitable for web-based automation and large-scale server integration. Given these technical constraints, AutoDock Vina the same docking engine employed internally by Chimera for its molecular docking calculations was selected. Its open-source accessibility, command-line operability, and compatibility with Python-based automation enabled seamless incorporation into the Flask backend [30]. Thus, the selection of AutoDock Vina reflects a balance between computational robustness, reproducibility, and automation feasibility, ensuring that Sanjevani delivers a reliable, literature-consistent docking experience while maintaining full accessibility through an interactive, browser-based interface.

**Table 12,13,14 :** Comparative benchmarking of molecular docking tools.

| S. No. | Ligand Name | Protein Target | Webina | CBDock2 | Chimera (kcal/mol) | PyRx (kcal/mol) | SwissDock |
| --- | --- | --- | --- | --- | --- | --- | --- |
| 1 | Epigallocatechin gallate | VEGFR2 | -8.5 | -8.5 | -7.6 | -8.8 | -7.7 |
| 2 | Curcumin | VEGFR2 + HER3 | -7.1<br>-7.7 | -9.7<br>-8.8 | -9.8<br>-8.2 | -6.9<br>-8.4 | -6.6<br>-5.3 |
| 3 | Genkwanin | VEGFR2 | -7.5 | -7.3 | -9.6 | -7.6 | -7.5 |
| 4 | Alpha-linolenic acid | VEGFR2 | -4.6 | -7.4 | -7.6 | -5.8 | -5.8 |
| 5 | Ferulic acid | VEGFR2 + HER2 | -6.2<br>-6.2 | -6.0<br>-6.0 | -7.4<br>-6.9 | -6.2<br>-5.7 | -6.2<br>-5.4 |
| 6 | Astragalin | h11 Beta-HSD1 | -9.0 | -9.7 | -8.9 | -9.0 | -9.4 |
| 7 | Carvacrol | h11 Beta-HSD1 | -5.7 | -5.8 | -5.8 | -5.7 | -5.7 |
| 8 | Cycloalivil | h11 Beta-HSD1 | -7.2 | -7.0 | -7.4 | -7.5 | -7.5 |
| 9 | Estra-1,3,5(10)-trien-17B-Ol | HCA-II | -7.1 | -7.6 | -7.5 | -7.5 | -7.5 |
| 10 | Pinoresinol | hCDC25 B | -7.2 | -7.3 | -7.2 | -7.3 | -6.9 |
| 11 | 8-Hydroxyacetophenone | HCA-II + h11 Beta-HSD1 | -5.6<br>-5.4 | -5.3<br>-5.5 | -5.7<br>-5.4 | -5.7<br>-5.7 | -5.8<br>-5.4 |
| 12 | Stigmastane | hCDC25 B | -7.6 | -7.0 | -7.5 | -7.5 | -7.1 |
| 13 | Allantoin | HSP90A A1 | -5.3 | -5.5 | -5.6 | -5.5 | -6.1 |

| Cell Color | Tools Representing | Color Codes | Representation |
| --- | --- | --- | --- |
| Light Grey | Chimera | Yellow | Matches with Literature |
| Light Blue | Webina | Blue | Matches with other literature |
| Light Orange | Pyrx | Green | Highest Affinity |
|  |  | Red | Closest to Literature Values |
|  |  | Pink | Matches with Literature along with highest value |

**Table 15:** Comparative performance evaluation of five molecular docking tools.

| Docking Tool | Matches with Literature | Matches with other literature/other docking tool result | Best Scores (Highest Affinity) | Closest to Literature Values | Strength |
| --- | --- | --- | --- | --- | --- |
| Chimera | 7 | 2 | 6 | 3 | Most reproducible & strong performer |
| Webina | 8 | 0 | 1 | 1 | Good reproducibility |
| PyRx | 1 | 2 | 4 | 6 | Very close to literature across many cases |
| SwissDock | 0 | 2 | 3 | 4 | Moderate consistency |
| CB-Dock2 | 0 | 1 | 5 | 3 | Good scoring ability but poor<br>reproducibility |

### AutoDock Vina Validation

The docking functionality of the platform was tested using the AutoDock Vina docking engine. A receptor-ligand pair from the Protein Data Bank structure 1A52 was used as a test case. The prepared receptor and ligand were successfully docked through the integrated AutoDock Vina pipeline, generating two binding poses with predicted binding affinities. The best binding mode showed an affinity score of −10.30 kcal/mol. The docking job completed successfully and the resulting protein–ligand complexes were visualized using the integrated molecular viewer. These results confirm that the docking workflow, execution of AutoDock Vina, and visualization of docking results, is functioning correctly within the platform.

**Fig 7:**
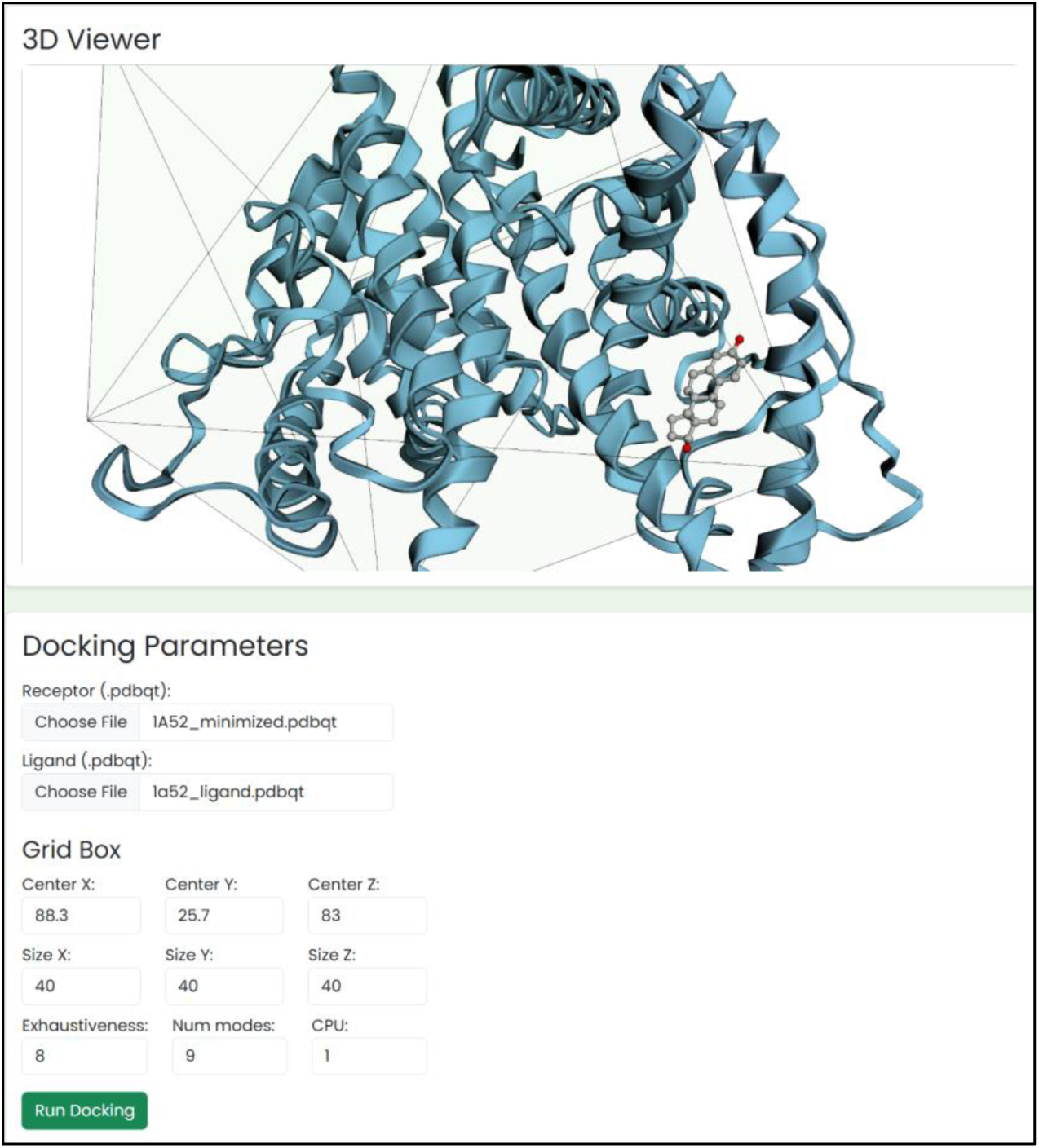
Docking 3D visualization and its Parameters.

**Fig 8:**
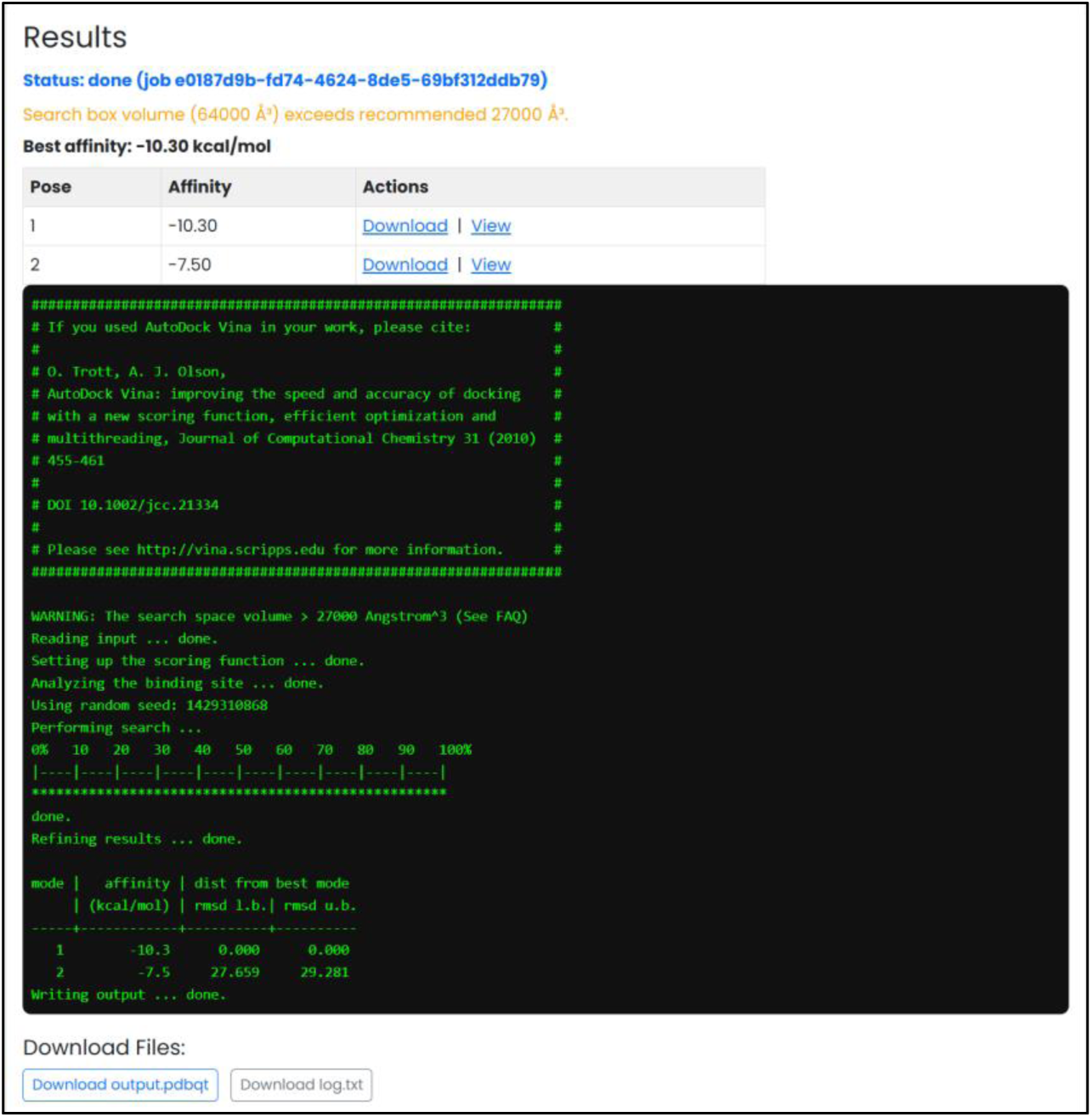
Results - Binding affinities of 1A52.

**Fig 9:**
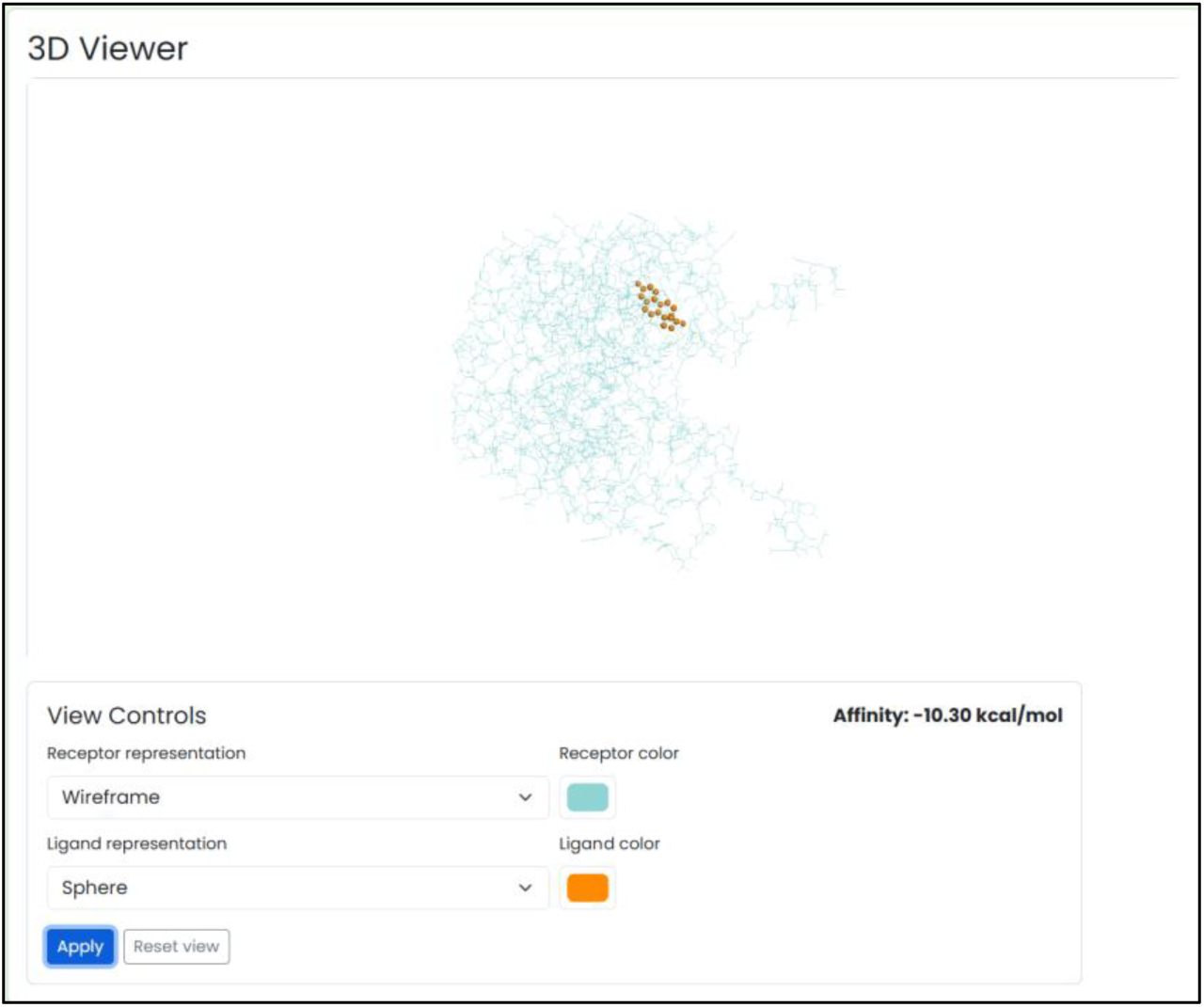
Visualization of highest affinity docking pose.

### ADMET-AI Validation

ADMET predictions for curcumin were obtained using the official admet.ai web service at admet.ai.greenstonebio.com and separately via our integrated admet.ai instance on the Sanjeevani server. ADMET predictions generated through the Sanjeevani-integrated ADMET-AI model were compared with predictions obtained from the official ADMET-AI web interface. The predicted values showed near-identical agreement, with only negligible differences (<10⁻⁶) attributable to floating-point precision and rounding differences between computational environments. The result is included in the **Supplementary table S2.**

## Discussion

The development of the Sanjevani database represents a significant advance in the field of computational phytochemistry and natural product based anticancer research. By integrating a curated library of 4,823 experimentally validated phytochemicals with machine learning, ADMET prediction and molecular docking modules, Sanjevani bridges the persistent gap between data collection, in silico bioactivity evaluation and ADMET based drug likeness assessment of the compounds. Compared with pre-existing repositories such as NPACT, CancerHSP, Dr. Duke’s Phytochemical and Ethnobotanical Database, ANPDB, AVPCD, BrNPDB and SANCDB, Sanjevani offers comprehensive data accessibility, downloadable 2D/3D structures, Molecular Docking and interactive prediction tools features and ADMET prediction that have historically been unavailable in most natural product databases. The high predictive accuracy (96.6%) achieved by the Support Vector Machine while Random Forest classifier achieved 95.8%, followed by K-Nearest Neighnors achieving 95.3% accuracy, underscores the robustness of combining ECFP4 fingerprints with physicochemical descriptors derived from RDKit. This hybrid representation effectively captures both substructural diversity and molecular property variance, enabling reliable discrimination between anticancer and inactive compounds. Importantly, the model’s balanced dataset created using the LIDeB decoy generator mitigated artificial enrichment, demonstrating strong generalization performance. These findings are consistent with recent reports emphasizing the efficacy of ML-models for virtual screening of phytochemicals. The integration of AutoDock Vina into the web platform further enhances the translational potential of Sanjevani. Users can now perform receptor–ligand docking directly through an interactive interface, eliminating the need for specialized software installation. This feature not only promotes reproducibility but also democratizes access to molecular modeling tools for researchers without computational expertise. Benchmarking experiments confirmed the reliability of the docking tool against published studies, validating its performance for binding affinity prediction across diverse protein targets. Despite these achievements, several challenges were encountered during development. Many natural product repositories such as NPCARE, and TIPdb originally included anticancer biological activity data; however, after their integration into the COCONUT framework, much of this information was not retained or displayed, resulting in a loss of explicit activity annotations. Similarly, databases like IMPPAT, though they catalog anticancer plants, do not always specify which phytochemicals are responsible for the observed activity, complicating data extraction and curation. Furthermore, assembling a negative dataset for machine learning was challenging due to the absence of experimentally proven non-anticancer phytochemicals, necessitating the generation of decoy molecules via the LIDeB tool. Finally, minor discrepancies were observed in docking affinity scores compared with literature reports, likely due to unreported variations in docking tool versions and parameter settings used across different studies. In summary, Sanjevani represents a comprehensive, integrative, and reproducible platform for anticancer phytochemical discovery. Although current limitations highlight areas for refinement, its continuous evolution driven by data expansion and advanced modelling will further strengthen its impact on computational oncology and natural product based drug research.

